# Histone demethylation and c-MYC activation enhance translational capacity in response to amino acid restriction

**DOI:** 10.1101/154419

**Authors:** Chen Cheng, Trent Su, Marco Morselli, Siavash K. Kurdistani

**Affiliations:** Department of Biological Chemistry, David Geffen School of Medicine, University of California, Los Angeles, California 90095, USA.; Department of Molecular, Cell and Developmental Biology, University of California, Los Angeles, California 90095, USA.; Department of Pathology and Laboratory Medicine, David Geffen School of Medicine, University of California, Los Angeles, California 90095, USA.; Eli and Edythe Broad Center of Regenerative Medicine and Stem Cell Research, David Geffen School of Medicine, University of California, Los Angeles, California 90095, USA.

**Author notes:** Correspondence to: Siavash Kurdistani.

## Abstract

Nutrient limitation may elicit adaptive epigenetic changes but the nature and mechanisms of the cellular response to specific nutrient deficiencies are incompletely understood. We report that depriving human cells of amino acids (AAs) induces specific loss of H4K20me1 from gene bodies and elevated binding of c-MYC at promoters genome-wide. These effects are most pronounced at ribosomal protein and translation initiation genes, which are upregulated, leading to enhanced protein synthetic capacity. Combination of H4K20 methyltransferase depletion and c-MYC over-expression in rich media is required and sufficient to recapitulate the effects of AA restriction. Our data reveal an unexpected and epigenetically implemented increase in translational capacity when AAs are limiting, likely to safeguard the proteome by making effective use of limited resources.

**One Sentence Summary:** Combination of H4K20me1 demethylation and c-MYC activation enhance translational capacity in response to amino acid restriction.

## Main Text

Cells respond to environmental challenges such as fluctuating nutrient levels partly through epigenetic mechanisms such as histone modifications and transcriptional regulation to reconfigure their metabolic or physiological states as adaptive measures (*1-3*). Understanding the nature and mechanisms underlying cellular response to nutrient availability can inform on fundamental regulatory processes and how they may be altered in human disease. From among the many nutrients that cells must sense and respond to, amino acids (AAs) are of particular interest because they regulate the mechanistic target of rapamycin complex 1 (mTORC1), a central hub for coordinating cell growth and metabolism (*4*). Although considerable progress has been made in recent years about how AAs activate mTORC1 and its downstream effects (*5*), much less is known about the epigenetic changes that may be required, especially independently of mTOR signaling, for effective adaptive response to limited AA availability.

We analyzed the effects of AA availability on histone modifications and discovered that deprivation of AAs induces genomewide loss of a single histone methylation site—histone H4 lysine 20 monomethylation (H4K20me1). HeLa cells were cultured for 16 hours in complete medium or media lacking DMEM components, which include vitamins, glucose, pyruvate, and amino acids. Removal of all 15 AAs, but none of the other media components, significantly reduced levels of H4K20me1 (Fig. 1A) without affecting the methylation levels of other lysine residues in histone H3 and H4 (Fig. S1A). The effect on H4K20me1 was not due to the absence of either the essential (EAA) or non-essential amino acids (NEAA) alone (Fig. 1B). In the absence of AAs, the cells remained viable over the course of the experiment (Fig. S1B) and recovered H4K20 mono-methylation when AAs were added back (Fig. 1C). We confirmed the effect of AA deprivation on H4K20me1 in HeLa cells with a different antibody (Fig. S1C) and observed the same effect in normal human bronchial/tracheal epithelial cells (HBTEC) and primary fetal lung fibroblasts (IMR90) (Fig. S1D).

**Figure 1.**
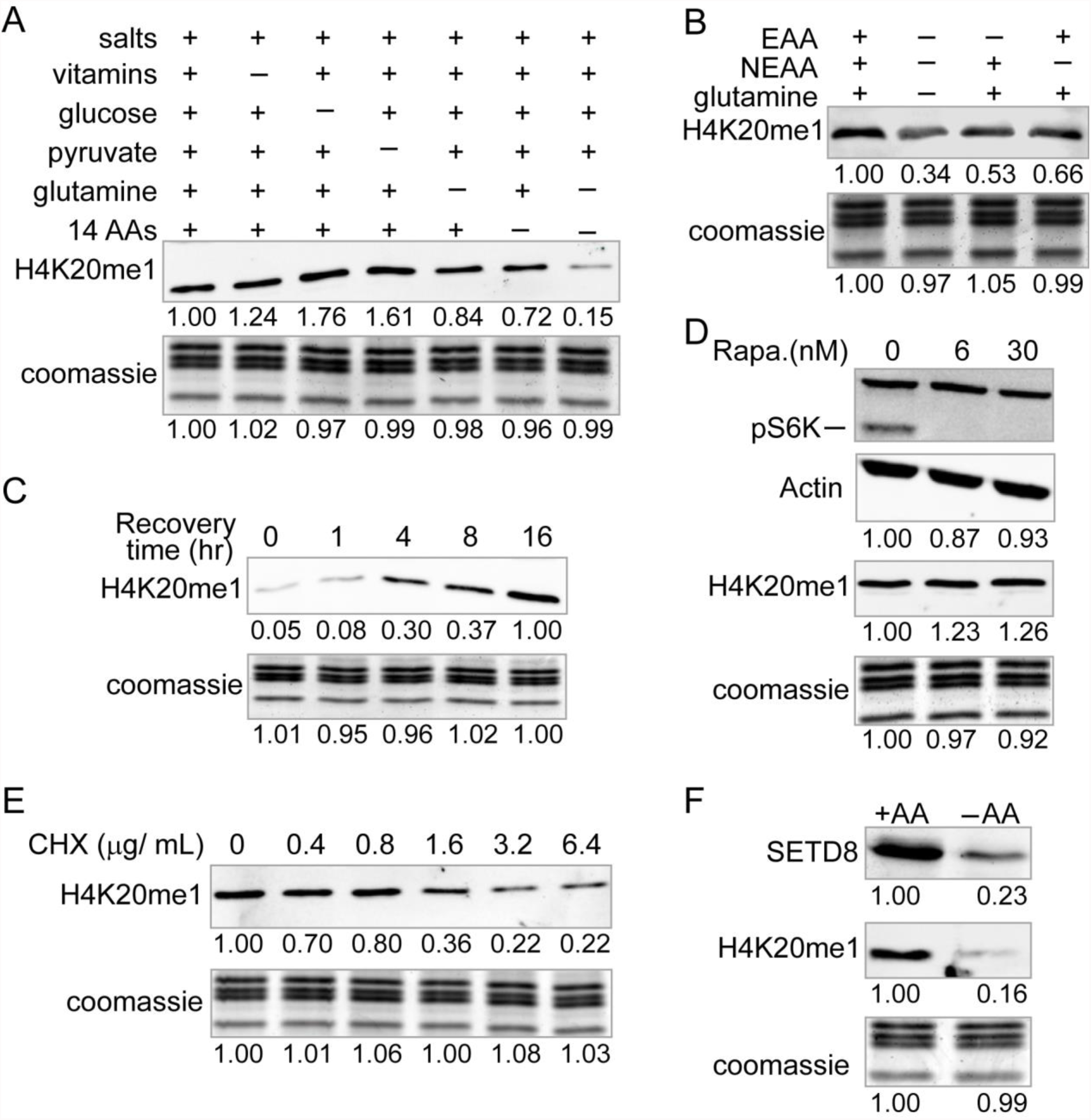
Global level of H4K20me1 varies proportionally to amino acid availability and translation capacity independent of mTOR signaling. (A-B) Quantitative Western blots (WB) of H4K20me1 in cells cultured for 16 hrs in the indicated media. 14 AA, non-glutamine amino acids in DMEM; EAA, essential amino acids; NEAA, non-essential amino acids. (C) WB of H4K20me1 in cells cultured without (−AA) amino acids for 16 hrs followed by recovery in complete media with AA for the indicated time. (D) WBs of phospho-S6K and H4K20me1 of cells treated with Rapamycin (Rapa.) in DMEM. (E) WB of H4K20me1 in cells treated with the indicated amount of cycloheximide (CHX) in DMEM. (F) WBs of H4K20me1 and SETD8 in cells cultured with (+AA) or without (−AA) amino acids.

As expected, removal of AAs inhibited mTOR activity as measured by its ability to phosphorylate S6K (Fig. S1E) (*6*). However, inhibiting mTOR activity with Rapamycin did not reduce the level of H4K20me1 (Fig. 1D), indicating that H4K20me1 is affected by AA availability in parallel to deactivation of mTOR signaling.

Since AA deprivation decreases protein production (*7*), we asked whether inhibition of translation in presence of AAs also affects H4K20me1. Indeed, addition of protein translation inhibitor cycloheximide (CHX) to complete DMEM caused a dose-dependent decrease in H4K20me1 levels (Fig. 1E). This indicates that the level of H4K20me1 is sensitive to productive translation and not AA availability *per se*.

The H4K20me1 methyltransferase SETD8 (Pr-SET7) level varies during cell cycle progression, with the lowest level in S phase and maximal level in late G2 (*8, 9*). We observed that the level of SETD8 decreased in cells cultured in media lacking AA (Fig. 1F). However, the decrease in SETD8 protein level was independent of cell cycle progression, since the cell cycle profiles of cells cultured with or without AAs for 16 hrs were comparable (Fig. S1F). Similarly, our treatment regimen with CHX, which resulted in a reduction of H4K20me1, had no appreciable effect on the cell cycle profile (Fig. S1G). Therefore, AA/translation-dependent change in H4K20me1 levels is associated with SETD8 turnover, and is not an indirect consequence of cell cycle changes.

To map the AA-dependent changes in global level of H4K20me1 to the genome, we performed chromatin immunoprecipitation-sequencing (ChIP-seq) analysis in cells cultured in presence or absence of AAs. H4K20me1 was enriched primarily in the gene bodies of 7589 genes (*10*), >95% of which exhibited decreased levels of H4K20me1 in the absence of AAs (Figs. 2A-2B). Consistent with the Western blot results (Fig. S1A), genome-wide distribution of H3K36me3, which is also enriched in the gene body, did not change substantially (Figs. S2A-S2B), confirming that AA restriction preferentially affects H4K20me1. The decrease in H4K20me1 did not appear to be due to increasing di- or tri-methylation of the same residue as the global levels of H4K20me2 and me3 were unaffected in the absence of AAs (Fig. S1A), and more importantly, H4K20me3 genome-wide enrichment pattern is physically distinct from H4K20me1 (Fig. S2C).

**Figure 2.**
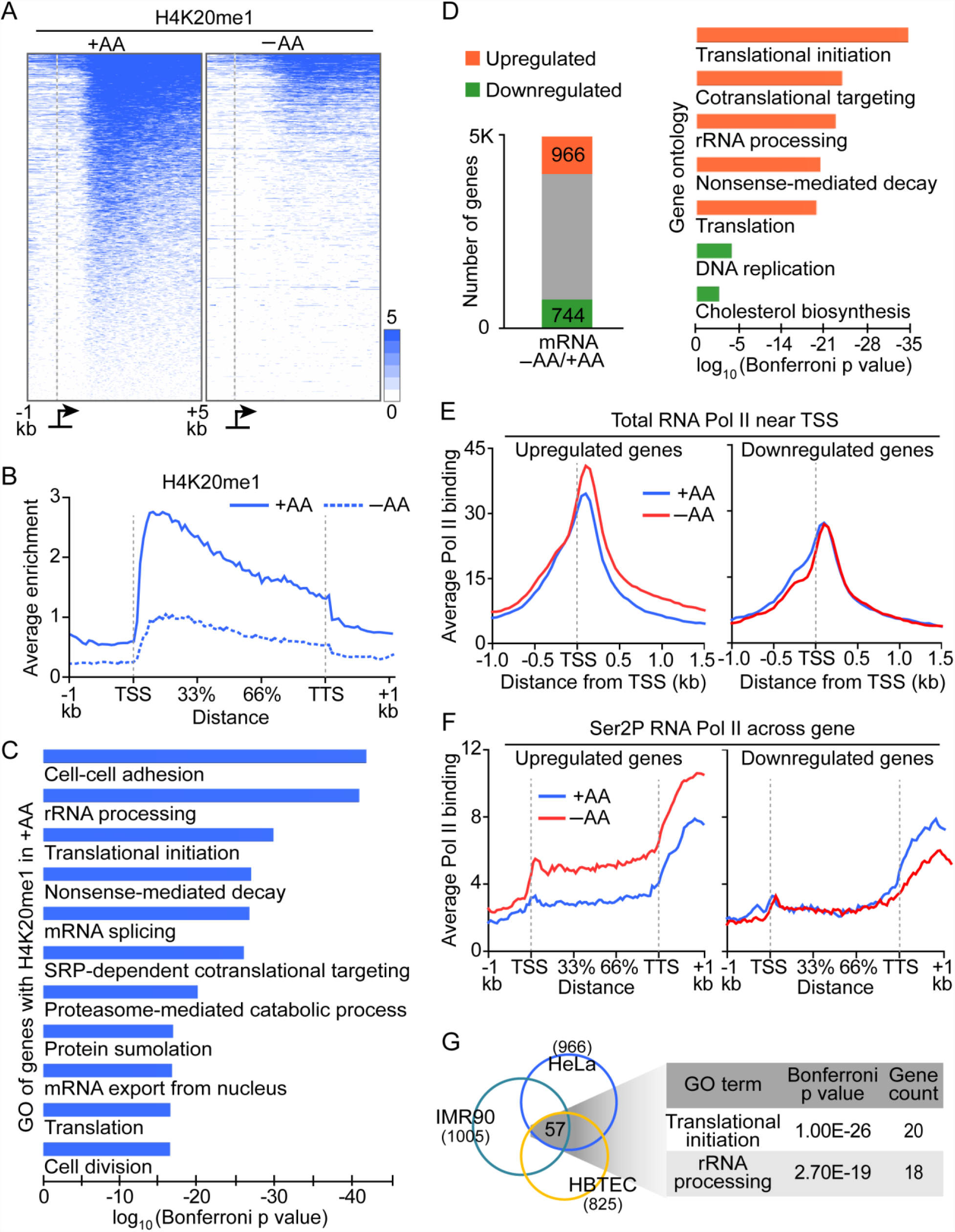
Amino acid restriction induces loss of H4K20me1 from gene bodies and upregulation of genes involved in translation. (A) Heat maps show the distribution of significant peaks of H4K20me1 in the indicated media for all genes with significant H4K20me1 enrichment in +AA. (B) Metagene plot of H4K20me1 levels for genes in (A) scaled to a distance of 3-kb between transcription start and termination sites. The distances away from the gene body are not scaled. (C) Gene ontology (GO) analysis of H4K20me1 associated genes from (A). (D) The number of upregulated or downregulated genes and their GO analysis in −AA. (E) Average significant peaks of total RNA Pol II, and (F) Metagene plots of significant Ser2P RNA Pol II peaks of the upregulated or downregulated genes. (G) The number of commonly upregulated genes and their GO analysis in −AA from the indicated cell types.

The most significantly enriched gene ontology (GO) terms for H4K20me1-associated genes include those involved in protein homeostasis (Fig. 2C). To correlate changes in H4K20me1 to gene expression, we performed mRNA-seq with a spike-in control in the same conditions as in ChIP-seq experiments. Compared to complete media, of all H4K20me1- associated genes, 966 and 744 were up- and down-regulated greater than 1.5-fold, respectively, in media lacking AAs (Fig. 2D, left panel). The genes within the upregulated group had the greatest loss of H4K20me1 (Fig. S2E) and were significantly enriched for genes involved in translation-related processes (Fig. 2D, right panel, orange bars). In contrast, the downregulated genes were only marginally enriched in specific GO terms (Fig. 2D, right panel, green bars). A closer analysis of the top GO terms among the upregulated genes (Fig. 2D) revealed a common set of genes encoding large and small ribosomal proteins (RPs), and translation initiation factors. A second biological replicate of mRNA-seq experiment, without a spike-in control, showed essentially identical functional enrichment of up- and down-regulated genes (Fig. S2D).

To determine whether differentially expressed genes were regulated at the level of transcription, we mapped the levels of total RNA polymerase II (Pol II) and elongating phosphor-Ser2 Pol II (Ser2P), as well as RNA-seq of the chromatin fraction (chromRNA-seq). The upregulated genes based on mRNA-seq had higher Pol II enrichment in both the promoter proximal region and gene body in the condition lacking AAs (Figs. 2E-2F, left panels). In contrast, there were no marked differences in Pol II occupancy between the two conditions among the downregulated genes (Figs. 2E-2F, right panels). Consistent with Pol II occupancy and mRNA-seq, chromRNA-seq showed increased nascent RNA associated with the upregulated but not with the downregulated genes (Figs. S2F-S2G). These data indicate that genes with increased mRNA levels because of AA deficiency are upregulated transcriptionally. The transcriptional response to AA restriction is not unique to HeLa cells as HBTEC and IMR90 cells also upregulated translation-related genes (Fig. S2H). In fact, the upregulated genes common to all three cell lines were highly significantly enriched with translational initiation and rRNA processing factors as well as RP genes (Fig. 2G).

We noticed that the promoter proximal sequences of upregulated translation-related genes contain the binding motif for transcription activator c-MYC and its binding partner MAX (Fig. S3A). The expression of c-MYC mRNA was also upregulated in the absence of AAs (Fig. S3B). We therefore examined the binding of c-MYC and found a substantial and genome-wide gain of c-MYC binding in the absence of AAs (Figs. 3A and S3C). Interestingly, there was a significantly greater increase in c-MYC binding within 1 kb of transcription start sites (TSS) of the H4K20me1 associated genes belonging to the GO terms “translational initiation” and “rRNA processing” which include RP genes (from Fig. 2C, n=217 genes) compared to randomly selected (100 permutations) sets of genes not in those terms (Figs. 3B and 3C). These observations indicate that in response to AA deprivation, a group of translation-related genes preferentially gain c-MYC, lose H4K20me1, and are upregulated.

**Figure 3.**
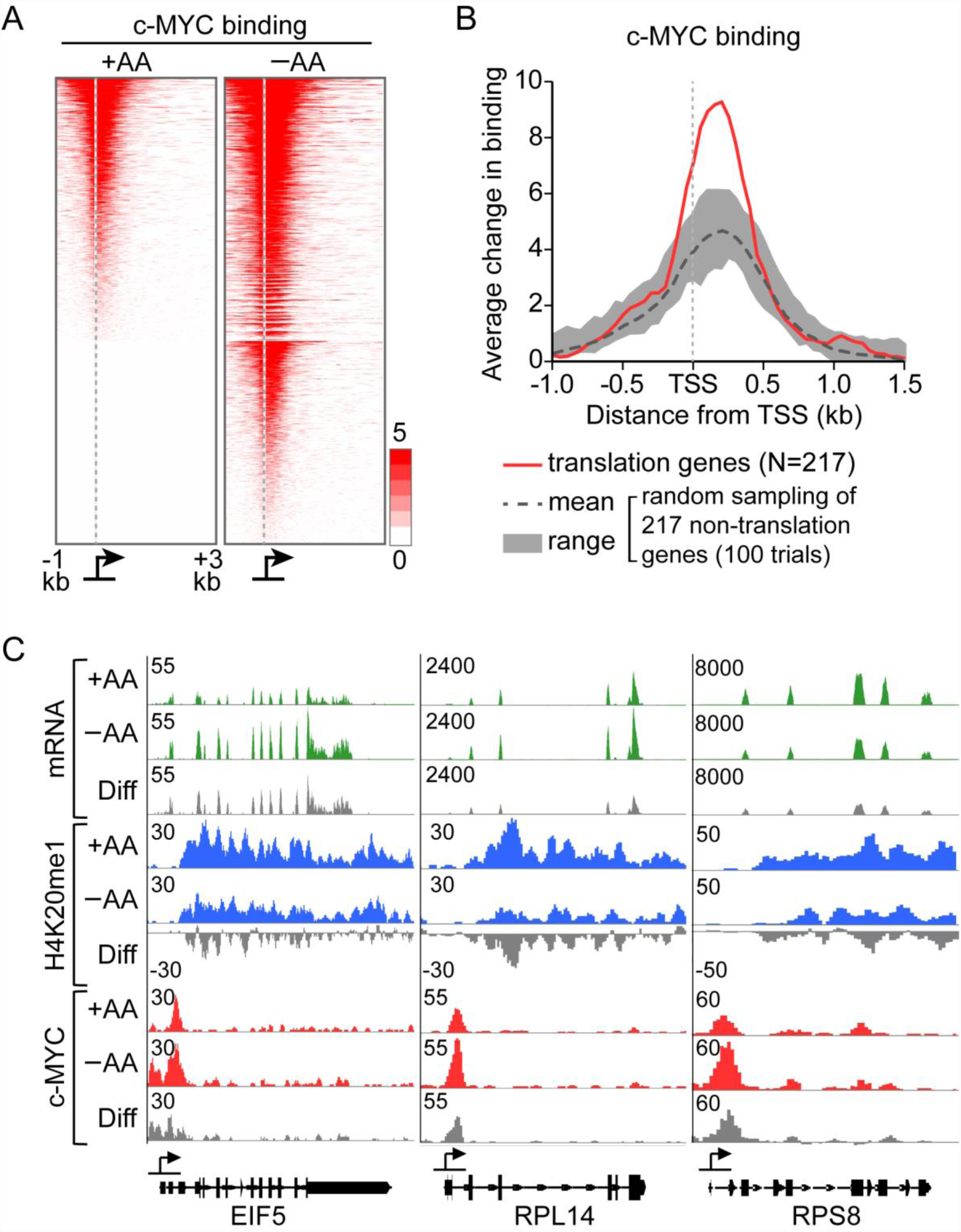
Amino acid restriction increases c-MYC binding genome-wide with preferential enrichment at genes involved in translation. (A) Heat maps show the distribution of significant peaks of c-MYC binding in the indicated media for all the genes with significant c-MYC binding enrichment within the indicated region in +AA or −AA. (B) Average differential binding of c-MYC around TSS of genes involved in translation (in red) or a group of randomly sampled non-translation related genes. The shaded grey and dashed line show the range and the mean of values from 100 sampling trials. (C) Genome browser tracks of representative translation initiation factor and large and small ribosomal protein genes.

To determine whether loss of H4K20me1 and gain of c-MYC binding are required to upregulate the expression of translation-related genes, we recapitulated these events in presence of AAs. We first knocked down *SETD8*, which effectively decreased H4K20me1 to a level similar to removal of AAs and which had no effect on c-MYC expression (Figs. 4A and S4A). Ectopic overexpression of c-MYC did not affect H4K20me1 level (Fig. 4B). Expression analyses indicated that none of the translation-related genes were upregulated by loss of SETD8 and H4K20me1, and only a small sub-set of translation initiation factors were upregulated by overexpression of c-MYC (Fig. 4C). However, concurrent knockdown of *SETD8* and overexpression of c-MYC resulted in increased expression of most initiation factors and RPs to the same extent as observed in the absence of AAs (Fig. 4C; see quantifications to the right of the heat map). Therefore, a combination of H4K20me1 loss and c-MYC gain is required and sufficient to upregulate the expression of translation-related genes in rich media.

**Figure 4.**
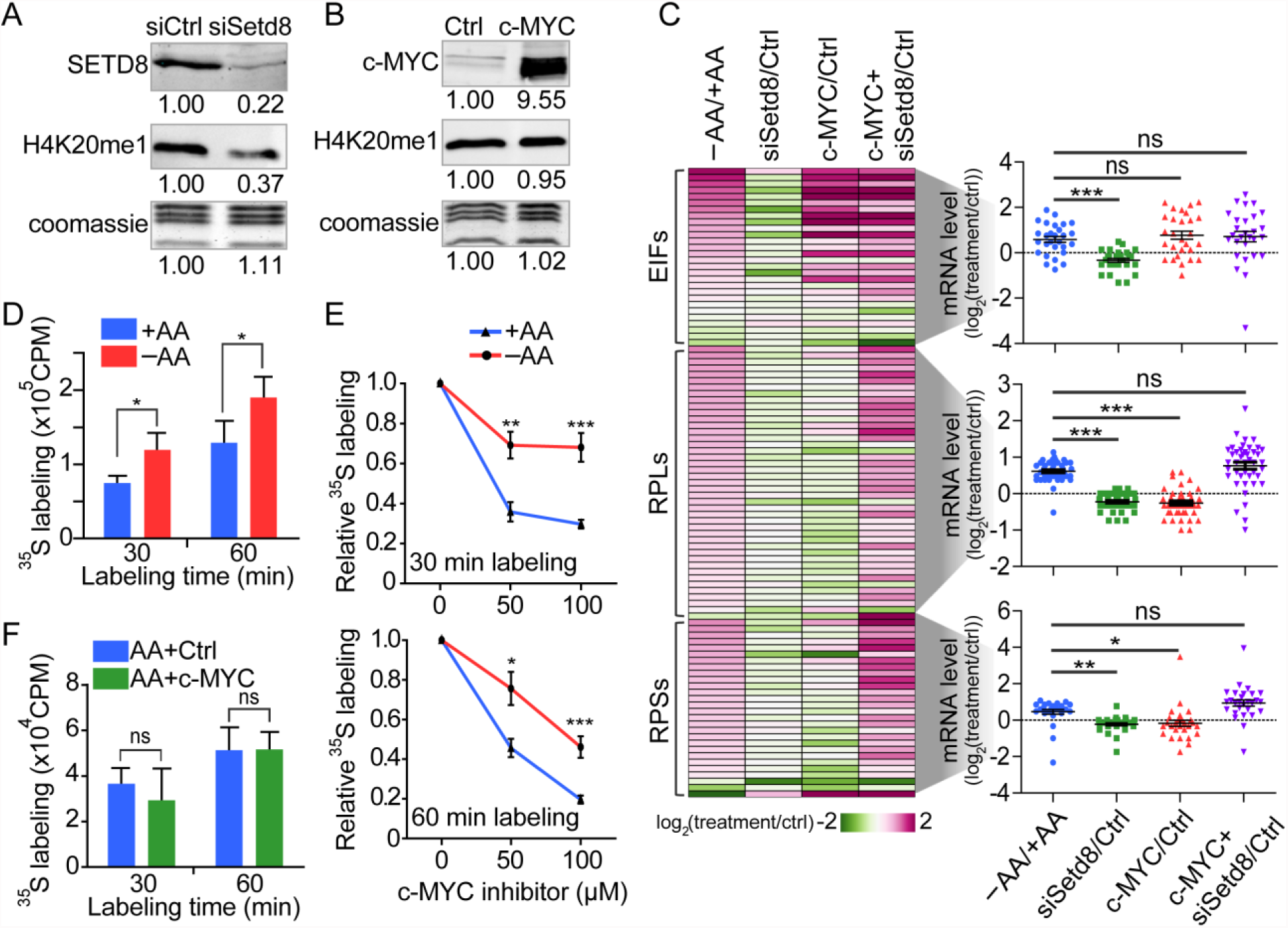
Concurrent loss of H4K20me1 and c-MYC overexpression are necessary for the upregulation of translational initiation and protein synthesis. (A) WBs of SETD8 and H4K20me1 in control and *SETD8* knockdown. (B) WBs of c-MYC and H4K20me1 in control and c-MYC overexpression. (C) Expression levels (heat map) and distributions (dot plots) of translation related genes in the indicated conditions. The t-test was used to calculate P values. ***P<0.0001, **P<0.001, *P<0.5, ns, not significant. (D) Levels of [^35^S]-methionine/cystiene incorporation by cells cultured with or without amino acids prior to pulse labeling. *P<0.01 (E) Relative levels of [^35^S]-methionine/cystiene incorporation by cells treated with DMSO or indicated concentrations of c-MYC inhibitor 10058-F4 prior to pulse labeling. *P<0.05, **P<0.01, *** P<0.001. (F) Levels of [^35^S]-methionine/cystiene incorporation by cells cultured in +AA and transfected with control or c-MYC expression plasmids for 48 hrs prior to pulse labeling.

Since depletion of amino acids from culture media reduces overall protein synthesis (*11*), we asked whether the upregulation of translation related genes is functionally relevant. Cells were cultured for 16 hours with or without AAs, and subjected to pulse-labeling in DMEM with or without AAs but supplemented with [^35^S]-labeled methionine and cysteine. Whole cell lysates were precipitated with trichloroacetic acid (TCA) to monitor the incorporation of radioactivity into total cellular proteins. After 30 or 60 minutes of labeling, cells cultured without AA prior to and during labeling showed significantly less incorporation as would be predicted by depletion of the cellular AA pool (Fig. S4B). Importantly, however, cells cultured without AAs prior to labeling but with full complement of AAs during labeling had significantly more incorporation of [^35^S]-labeled methionine/cysteine than those continuously cultured in complete media (Fig. 4D). These observations indicate that when AAs become limiting, a subset of RP and translation initiation genes are upregulated and translation capacity is increased.

The baseline translation capacity in rich media and its elevated levels in response to the absence of amino acids were dependent partly on the function of c-MYC. Cells cultured in presence of the c-MYC inhibitor 10058-F4, which antagonizes c-MYC binding to DNA (*12*), for 8 hours prior to [^35^S] labeling, had significantly lower protein synthesis (Fig. 4E). Interestingly, the c-MYC inhibitor was more effective at inhibiting protein synthesis when cells were cultured in the presence of AAs than in their absence (Fig. 4E). This is consistent with increased expression and binding of c-MYC under AA deprivation condition (Fig. 3), which may render the inhibitor drug less effective at inactivating c-MYC. However, just as overexpression of c-MYC alone was insufficient to fully induce the expression of translation-related genes (Fig. 4C), it also did not increase protein synthetic capacity in rich media (Fig. 4F). We therefore conclude that c-MYC and H4K20me1 coordinately regulate the expression of a subset of translation-related genes, modulating translational capacity in response to availability of AAs or productive translation.

The function of c-MYC in transcriptional regulation of ribosome biogenesis has been well characterized (*13*). Our data has now uncovered the importance of H4K20me1 for the ability of c-MYC to further enhance translational capacity in response to AA deprivation. H4K20me1 has been implicated in both gene activation and repression (*10*) but in our condition, H4K20me1 appears to play an inhibitory role for translation-related genes. Since there are no confirmed demethylases for H4K20me1, whether its loss in the absence of AAs is due to passive or active demethylation or a combination of both remains an open question. H4K20me1 has a half-life of ~9 hrs (*14*), suggesting that in principle some loss of methylation could occur passively in the course of our experiments as SETD8 levels fall. Nevertheless, the parallel functions of c-MYC and chromatin to coordinate a counterintuitive increase in translational capacity when AAs are limiting could prepare the cell for faster recovery when AAs are replenished. This notion may also have implications for understanding how c-MYC drives cancer metabolism and progression, especially in the context of enhancing protein biosynthesis in cancer (*15*).

## Acknowledgments

We thank Oscar A. Campos, Narsis Attar and Maria Vogelauer for critical review of the manuscript, and acknowledge the services of the UCLA Broad Stem Cell Center Sequencing Core. **Funding:** This work was supported by the NIH grant CA178415 to S.K.K. **Data and materials availability:** All the sequencing data generated in this study have been deposited to the Gene Expression Omnibus database under the accession number GSE100304.

**Supplementary Materials:**

Materials and Methods Table S1

Fig S1 – S4

References (16 – 29)

## Supplementary Materials

## Materials and Methods

### Cell Culture

HeLa and IMR90 cells were maintained in DMEM (Cellgro #10-013-CV) supplemented with 10% fetal bovine serum (FBS) (Thermo Fisher HyClone #SV30014.03). HBTEC cells were obtained from Cell Systems and maintained in the recommended medium (Cell Systems #FC-0035). For all experiments with custom media (detailed below), the cells were treated for 16 hrs unless otherwise indicated.

### Media Preparation

Custom media were prepared where applicable as indicated in the figures and results. Media containing complete amino acids (AA) were made from DMEM powder (US Biological #D9802) and supplemented with an additional 3.5 g/L D-glucose (for a total of 4.5 g/L), 1 mM sodium pyruvate (Gibco #11360070), 3.7 g/L sodium bicarbonate, and 0.0159 g/L sodium phenol red (US Biological #P4040). Media without AA were made from DMEM without amino acids powder (US Biological #D9800-013) and supplemented with an additional 3.5 g/L D-glucose, 1 mM sodium pyruvate (Gibco #11360070), and 3.7 g/L sodium bicarbonate. pH of media was adjusted to 7.2 by the addition of HCl before being filter sterilized and supplemented with 10% dialyzed FBS (Gibco #26400-044). These +AA and −AA media were used for all experiments comparing the two conditions.

Custom dropout media were made by combining individual components of DMEM from sources as indicated below. A 10X inorganic salt mixture was made according to Earle’s balanced salts recipe (Sigma #E7510). D-Glucose was supplemented at 4.5 g/L. Individual amino acids (except glutamine) were made at 100X of the concentrations found in DMEM. Premade solutions of vitamins mixture (Sigma #M6895), sodium pyruvate (Gibco #11360070), and L-glutamine (Gibco #25030081) were used. All dropout media were supplemented with 44 mM sodium bicarbonate. pH was adjusted to 7.2 before being filter sterilized and supplemented with 10% dialyzed FBS (Gibco #26400-044). Media were conditioned at 37°C and 5% CO2 for 6 hrs to overnight before adding to cells.

### Acid Extraction of Histones and Whole Cell Extracts

Acid extracted histones were prepared as described previously (*16*). Briefly, nuclei were collected by lysing 70-80% confluent cells in a 10-cm plate with a hypotonic buffer (10 mM HEPES pH 7.9, 1.5 mM MgCl_2_, 10 mM KCl, 340 mM sucrose, 10% glycerol, 1 mM DTT, protease inhibitors) and then acid-extracted with H_2_SO_4_. The proteins were collected with trichloroacetic acid (TCA) precipitation and dissolved in 100 μL water for concentration measurement. For whole cell extracts (WCE), cells were pelleted after PBS washing and resuspended in Laemmli buffer (2% SDS, 10% glycerol, 60 mM Tris-HCl pH 6.8). The extract was boiled for 10 min and sheared by passing through a 0.7 mm needle (BD #305156) 10 times.

### Subcellular Fractionation

Chromatin fraction was isolated as described previously (*17*). Briefly, nuclei were collected the same way as for acid histone extraction as described above and resuspended in lysis buffer B (3 mM EDTA, 0.2 mM EGTA, 1 mM DTT, protease inhibitors). The nuclear extract was pelleted and resuspended in SDS sample buffer for WB analysis.

### Coomassie Gel and Western Blotting (WB) Analyses

Analysis and quantification of protein gels and WB were performed using the Odyssey Infrared Imaging System (LI-COR Biosciences). Histone protein concentration was initially quantified with the BCA assay (Thermo Scientific #23225) and then normalized between samples by quantifying the four histone bands on gels stained with SimplyBlue (Invitrogen #LC6060, referred as coomassie in figures). After normalizing the concentration, 1 μg of protein was loaded in duplicate gels for WB and a second gel for SimplyBlue staining to ensure equal loading. This step—equal loading of histone based on coomassie staining—is critical for accurate determination of histone modification levels. For WCE and chromatin fraction, samples were initially loaded in equal volumes on a gel stained with SimplyBlue to approximate relative amounts loaded. The loading volume was normalized between samples by quantifying the four histone bands and a region corresponding to the sizes of the proteins to be analyzed in subsequent WB. WBs were performed on acid-extracted histones for histone modifications, on WCE for SETD8, and on chromatin fraction for c-MYC. WBs were performed with 15% polyacrylamide gels for histone modifications and with 10% gels for non-histone proteins. The gels were transferred onto Immobilon-FL PVDF membrane (Millipore #IPFL00010). Primary and secondary antibodies and dilutions used are listed in Table S1.

### mTOR Inhibition with Rapamycin

The 5 mM Rapamycin (Calbiochem #553211) stock provided by vendor was diluted to a 6 μM working solution with DMSO and added to DMEM at final concentration of 6 or 30 nM. Control media were prepared by diluting DMSO into media in the same amount used in the treatment. Cells were treated in Rapamycin or DMSO containing media for 8 hrs.

### Protein Synthesis Inhibition with Cycloheximide

A stock solution of cycloheximide (Sigma #C4859) was diluted to a 1.6 mg/mL working solution with DMSO and added to DMEM at final concentrations ranging from 0.4 to 6.4 μg/mL. Control media were prepared by diluting DMSO into media in the same amount as in the highest amount of treatment. Cells were treated in cycloheximide or DMSO containing media for 16 hrs.

### Flowcytometry Analysis

Cells were washed with cold PBS immediately following treatment and dissociated into single cells with trypsin. 1x10^6^ cells were fixed in ethanol and stained with propidium iodide (Invitrogen #P3566) for flow cytometry analysis using the FACSCalibur system (BD Biosciences). DNA content and cell cycle profile were analyzed with the ModFit LT software (Verity Software House).

### Cell viability

Cells were dissociated and stained with Trypan Blue solution (Biorad, #145-0021). Cell viability was assessed using the TC10™ automated cell counter (Biorad, #145-0010).

### c-MYC Inhibition with 10058F4

c-MYC inhibitor 10058F4 was dissolved in DMSO to make a 25 mM stock solution and diluted into +AA or −AA media at a final concentration of 50 or 100 μM. Control media were prepared by diluting DMSO into media at a concentration of 4 μL of DMSO per mL of medium. Cells were cultured for 8 hrs in drug free media followed by 8 additional hrs in media containing the inhibitor or DMSO.

### Chromatin Immunoprecipitation (ChIP)

H4K20me1 and H3K36me3 ChIP were performed as described previously (*18*). Specifically, after being cultured for 16 hrs in +AA or −AA media, HeLa cells were cross-linked in 1% formaldehyde for 10 min at 37°C and then neutralized with 140 mM glycine for 5 min. Cross-linked cells were scraped from the plates and washed with PBS containing protease inhibitors (Roche #11836145001). 2x10^7^ cells were resuspended in 400 μL of ChIP lysis buffer (1% SDS, 50 mM Tris-HCl pH 8, 20 mM EDTA, protease inhibitors) and incubated for 10 min on ice. Immediately, lysates were sonicated using a Misonix sonicator. Sheared lysates were diluted with dilution buffer (16.7 mM Tris-HCl pH 8, 0.01% SDS, 1.1% Triton X-100, 1.2 mM EDTA, 167 mM NaCl) and pre-cleared with Protein G beads (ThermoFisher #10003D) for 2 hrs before immunoprecipitation (IP). 5% of pre-cleared lysate was saved as input. 100 μL (approximately 5x10^6^ cells) of the lysate was incubated overnight with a given antibody (Table 1). Chromatin-antibody complexes were captured with Protein G beads for 2 hrs. Beads were washed twice each with low salt wash buffer A (140 mM NaCl, 50 mM HEPES pH 7.9, 0.1% SDS, 1% Triton X-100, 0.1% Na-deoxycholate), high salt buffer B (500 mM NaCl, 50 mM HEPES pH 7.9, 0.1% SDS, 1% Triton X-100, 0.1% Na-deoxycholate), LiCl buffer (20 mM Tris-HCl pH 8, 250 mM LiCl, 1 mM EDTA, 0.5% Na-deoxycholate, 0.5% NP-40), and with 1x TE. Protein-DNA complexes were eluted from beads at 65°C once each with 100 μL elution buffer (50 mM Tris-HCl pH 8, 1 mM EDTA) and with 150 μL TE containing 0.67% SDS. Eluates were incubated overnight at 65°C to reverse the cross-links and treated with RNase A and Proteinase K. DNA was subsequently extracted using phenol:chloroform:isoamyl alcohol (Invitrogen #15593031).

RNA Pol II and c-MYC ChIP experiments were performed using the same procedure as histone methylation ChIP with the following modifications. Lysates for c-MYC ChIP were sonicated with Bioruptor, and 2x10^7^ cells were used for each IP. Cells for RNA Pol II ChIP were resuspended and sonicated in buffers described in (*19*). Briefly, 6x10^6^ cross-linked cells were resuspended, incubated for 10 min, and then pelleted once each in 600 μL of lysis buffer 1 (50 mM HEPES-KOH pH 7.5, 140 mM NaCl, 1 mM EDTA, 10% glycerol, 0.5% NP-40, 0.25% Triton X-100, protease inhibitors) at 4°C and in 600 μL of lysis buffer 2 (10 mM Tris-HCl pH 8, 200 mM NaCl, 1 mM EDTA, 0.5 mM EGTA, protease inhibitors) at room temperature. Cells were then sonicated in 200 μL of lysis buffer 3 (10 mM Tris-HCl pH 8, 200 mM NaCl, 1 mM EDTA, 0.5 mM EGTA, 0.1% Na-deoxycholate, 0.5% *N*-lauroylsarcosine, protease inhibitors).

### ChIP Sequencing Library Preparation

DNA obtained from ChIP experiments was quantified using Qubit assays (Invitrogen #Q32854). 2 ng DNA from each ChIP and corresponding input samples were used to prepare libraries using the NuGEN Ovation Ultralow Library System (NuGEN Technologies #0331) or the KAPA LTP kit (KAPA Biosystems #8230) and sequenced with Illumina Hi-seq platforms to obtain 50 bp-long reads.

### ChIP-seq Data Analysis

Reads were aligned to the Human genome (hg19) using Bowtie (*20*) with parameters that allow up to 2-bp mismatch (-v2) and ensure only unique aligning reads were collected (-m1). For all aligned data files, duplicated reads were removed using Samtools (*21*). To estimate ChIP enrichment, IP and corresponding input files were randomly sub-sampled to equalize the number of reads. ChIP enrichment was determined using an algorithm described previously (*18*). Briefly, the software allows user to define a window size to tile the genome and a Poisson distribution p-value cutoff to call a window significant. For all experiments reported in this study, the genome was tiled into 50-bp windows. Significant peaks are defined as those windows with Posisson p< 10^−3^ and the same p value cutoff is met in the two neighboring windows.

### Calculation of ChIP Enrichment over Specific Genomic Features

The enrichment profiles for significant peaks near TSS or across entire genes were generated using Cis-regulatory Elements Annotation System (CEAS) (*22*). H4K20me1 associated genes (Figs. 2A-2B) are those with significant H4K20me1 enrichment within the region -1 to +5 kb around the transcription start site (TSS). The enrichments profiles were then used to graph average profiles around TSS and Metagene or to generate heat maps.

### mRNA-seq and chromatin RNA-seq Library Preparation

For mRNA-seq, total RNA was extracted using the Trizol reagents (Ambion #15596026) and treated with TURBO DNase (Ambion #AM2239). 1 μg of total RNA was used to prepare sequencing library using the Illumina TruSeq RNA sample preparation kit. A mixture of external RNA spike-in controls (Ambion #4456740) was added to total RNA prior to library preparation (*23*). Libraries were sequenced with Illumina Hi-seq platforms to obtain 50 bp-long reads. No spike-in control was added to experiment 2 (Fig. S2D), and 100 bp-long reads were obtained from those two libraries.

For chromatin RNA-seq, subcellular fractionation and sequencing libraries were prepared as described previously (*24*). Libraries were sequenced with Illumina Hi-seq platforms to obtain 50 bp-long reads.

### RNA-seq Data Analysis

Reads were mapped to the Human genome (hg19) and to spike-in reference sequences using default parameters of TopHat (*25*). SAMMate software (*26*) was used to determine the transcript RPKM (reads per kilobase of exon per million of reads) for mRNA-seq libraries. The sums of RPKM values from the spike-in controls were used to normalize across samples and to adjust transcripts RPKM.

### Gene ontology analysis

Gene ontology analysis was performed by uploading Genebank accession numbers for gene lists of interest to DAVID Bioinformatics Resources 6.8 (*27, 28*). Significant GO terms were defined as those terms with a minimal of 10 genes and with a Bonferroni corrected p value< 0.01.

### Motif enrichment analysis

Enriched motifs were identified using the HOMER software (*29*). The perl script <u>findMotifs.pl</u> was used to find motifs enriched within the region -500 to +750 bp from the TSS of a list of genes.

### siRNA and c-MYC plasmid transfection

SETD8 (NM_020382) knockdown was performed using a predesigned Dicer-substrate siRNA (DsiRNA) targeting exon 8/UTR of SETD8 (IDT DNA #HSC.RNAI.N020382.12.4). HeLa cells were transfected with 50 nM of DsiRNA using Lipofectamine RNAiMAX (Invitrogen #13778) for 48 to 72 hrs. Control samples were transfected with 50 nM of scrambled negative control DsiRNA (IDT DNA #51-01-19-09). c-MYC was expressed by a pcDNA3-cmyc plasmid (addgene #16011). HeLa cells in 10-cm plates were transfected with 10 μg of DNA using BioT reagent (Bioland Scientific #B01) for 24 to 48 hrs.

### Measuring Protein Synthesis by Pulse Labeling

After being cultured with the indicated conditions, cells were washed and incubated in DMEM lacking methionine and cysteine for 15 min at 37°C. The media were replaced with the same DMEM supplemented with 0.1 mCi/mL [^35^S]-Methionine-Cysteine (PerkinElmer NEG77200) for 30 or 60 min. After radioactive labeling, cells were washed twice with cold PBS and collected in 0.5 mL PBS. The labeled cells were resuspended in IP lysis buffer (250 mM NaCl, 50 mM Tris-HCl pH8, 5 mM EDTA, 0.5% NP-40, protease inhibitors), incubated on ice for 30 min, and centrifuged for 5 min at 10000x g. The supernatants were quantified using Qubit protein assay (Invitrogen #Q33212). Equal amount of lysates, as determined by protein concentrations, were precipitated with TCA using BSA as a carrier protein. The precipitated proteins were filtered onto 2.5-cm glass microfiber filter disks (Whatman #1822-025) and washed twice each with ice cold 10% TCA and 100% ethanol. The filters were air dried for 30 min and placed into vials containing scintillation fluid (BD #SX18). [^35^S]-activity was measured by a liquid scintillation analyzer (PerkinElmer Tri-Carb 2800TR).

**Table S1.** Primary and Secondary Antibodies used in WB and ChIP

| Antibodies | Source | Identifier | Dilution |
| --- | --- | --- | --- |
| H4K20me1 | Abcam | Ab9051 | WB: 1/4000<br>ChIP: 5 µg |
| H4K20me1 #2 (Fig. S1C) | Active Motif | 39175 | WB: 1/2000 |
| H3K4me2 | Abcam | Ab32356 | WB: 1/2000 |
| H3K4me3 | Abcam | Ab8580 | WB: 1 µg/mL |
| H3K9me1 | Active Motif | 39681 | WB: 1/500 |
| H3K9me2 | Abcam | Ab1220 | WB: 1/1000 |
| H3K36me1 | Abcam | Ab9048 | WB: 1/1000 |
| H3K36me3 | Abcam | Ab9050 | WB: 1 µg/mL |
| H3K79me2 | Grunstein Lab | #532 | WB: 1/3000 |
| H4K20me2 | Abcam | Ab9052 | WB: 1/1000 |
| H4K20me3 | Abcam | Ab9053 | WB: 1/1000 |
| Phospho-p70 S6 kinase (Thr389) | Cell Signaling | 9205 | WB: 1/1000 |
| Actin | Santa Cruz | Sc-8432 | WB: 1/2000 |
| SETD8 (hPR-SET7) | Millipore | 06-1304 | WB: 1/500 |
| c-MYC | Santa Cruz | sc-764 | WB: 1/2000<br>ChIP: 10 µg |
| RNA Pol II (8WG16) | Covance | MMS-126R | WB: ChIP: 1/50 |
| PhosphoSer2-RNA Pol II CTD | Abcam | Ab5095 | WB: ChIP: 5 µg |
| IRDye® 800CW Goat anti-Rabbit IgG | LI-COR | 926-32211 | WB: 1/10000 |
| IRDye® 800CW Goat anti-Mouse IgG | LI-COR | 926-32210 | Wb: 1/10000 |

**Figure S1.**
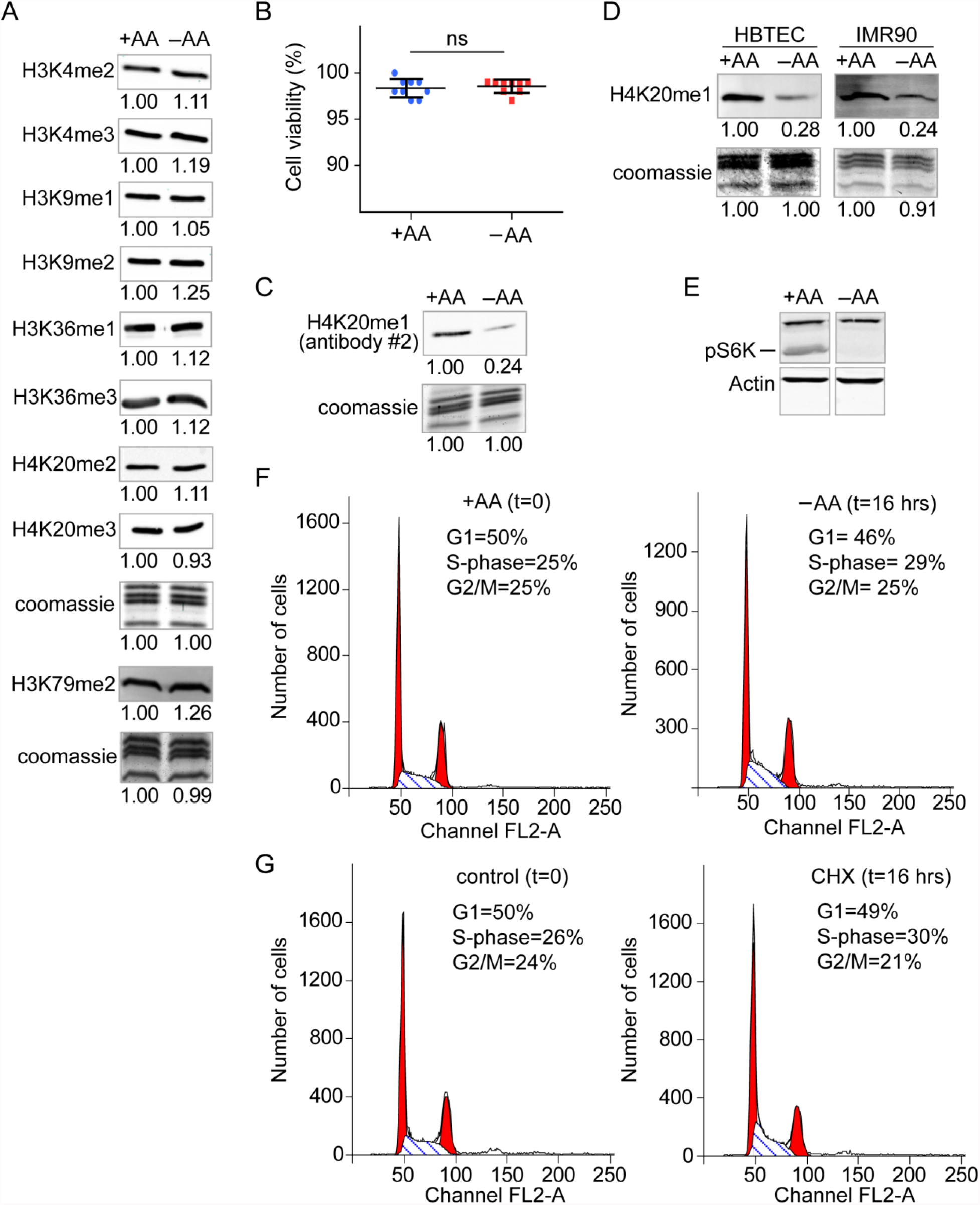
(A) WBs of histone methylation of cells cultured with (+AA) or without (−AA) amino acids for 16 hrs. The same preparation of histones was used to perform all WBs in this figure except for H3K79me2, which was done at a later time with its own loading control (bottom panel). (B) Viability of cells cultured in +AA or −AA using Trypan Blue staining. (C) WBs of H4K20me1 using a second antibody (Active Motif cat. no. 39175; lot no. 01008001) in HeLa cells cultured with (+AA) or without (-AA) amino acids. (Note that the Abcam H4K20me1 antibody (cat. no. 9051; lot no. GR79450-1) was used for all other WB and ChIP experiments in this study.) (D) WBs of H4K20me1 in HBTEC or IMR90 cells cultured with (+AA) or without (-AA) amino acids. (E) WBs of phospho-S6K of cells cultured in +AA or −AA. (F-G) Flow cytometry analysis of propidium iodine (PI) stained HeLa cells cultured in the indicated conditions. The percentage of cells in each phase of the cell cycle is indicated.

**Figure S2.**
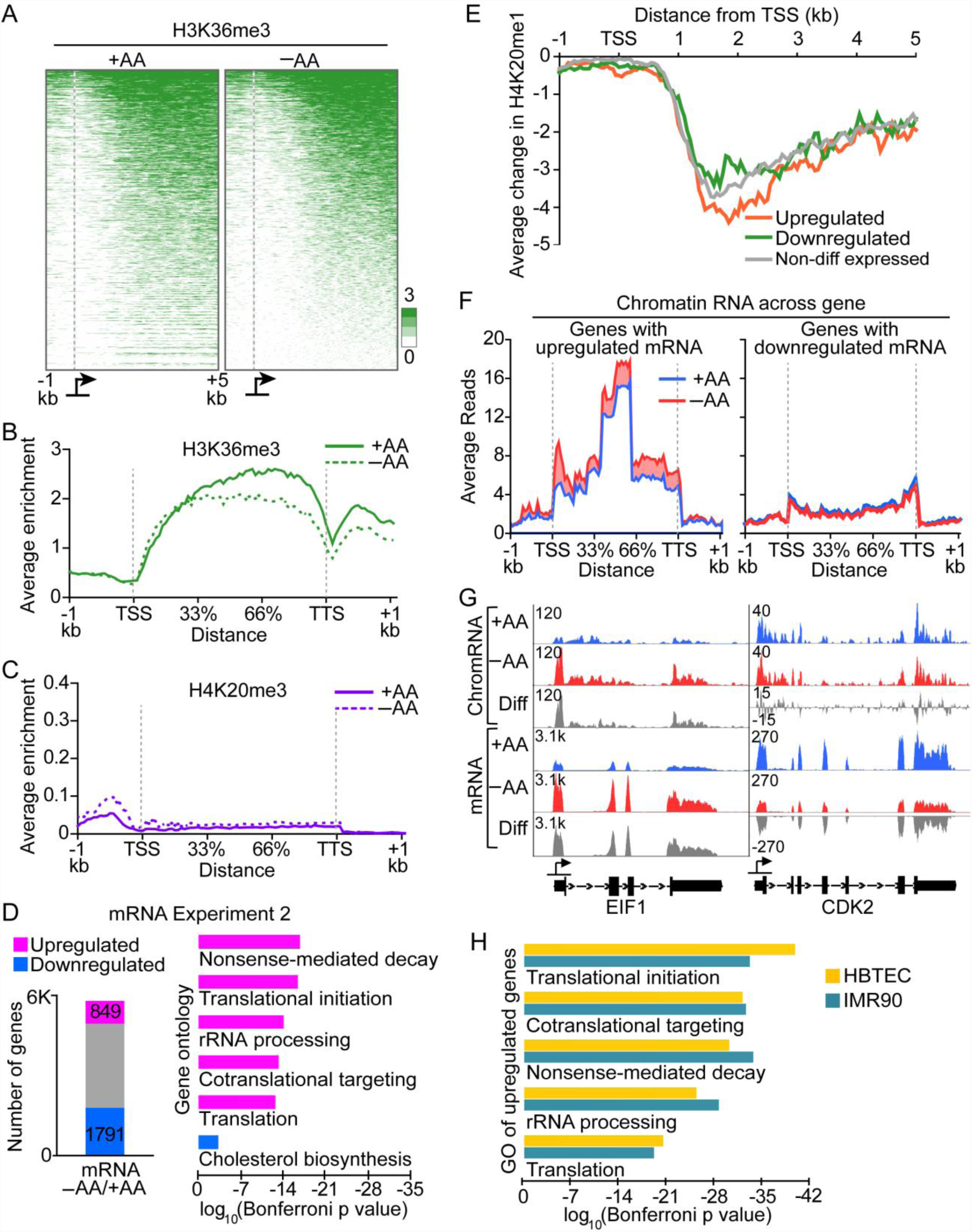
(A) Heat maps showing the distribution of significant peaks of H3K36me3 in the indicated media within the region -1 to +5 kb around the transcription start site (TSS). Included in the heat maps are all the genes with significant H3K36me3 enrichment within the 6-kb region in +AA. (B-C) Metagene plot showing the significant H3K36me3 peaks of the same group of genes shown in figure S2A. (D) Related to Fig. 2D. Biological replicates of mRNA-seq of cells cultured in +AA or −AA without a spike-in control. Left panel, numbers of genes with mRNA upregulated or downregulated greater than 1.5-fold in −AA compared to +AA. Right panel, GO analysis of upregulated or downregulated genes. (E) Average differential enrichment of H4K20me1 (−AA vs. +AA) within the region -1 to +5 kb around TSS from upregulated, downregulated, or non-differentially expressed genes (as identified in figure 2D). (F) Metagene plot showing chromatin fraction RNA-seq (chromRNA) coverage of the upregulated or downregulated group of genes shown in figure 2D. (G) Genome browser tracks showing chromRNA and mRNA coverage in +AA and −AA from representative upregulated or downregulated genes shown in figure 2D-F. (H) GO analysis of upregulated genes in −AA vs. +AA from normal primary IMR90 and HBTEC cells.

**Figure S3.**
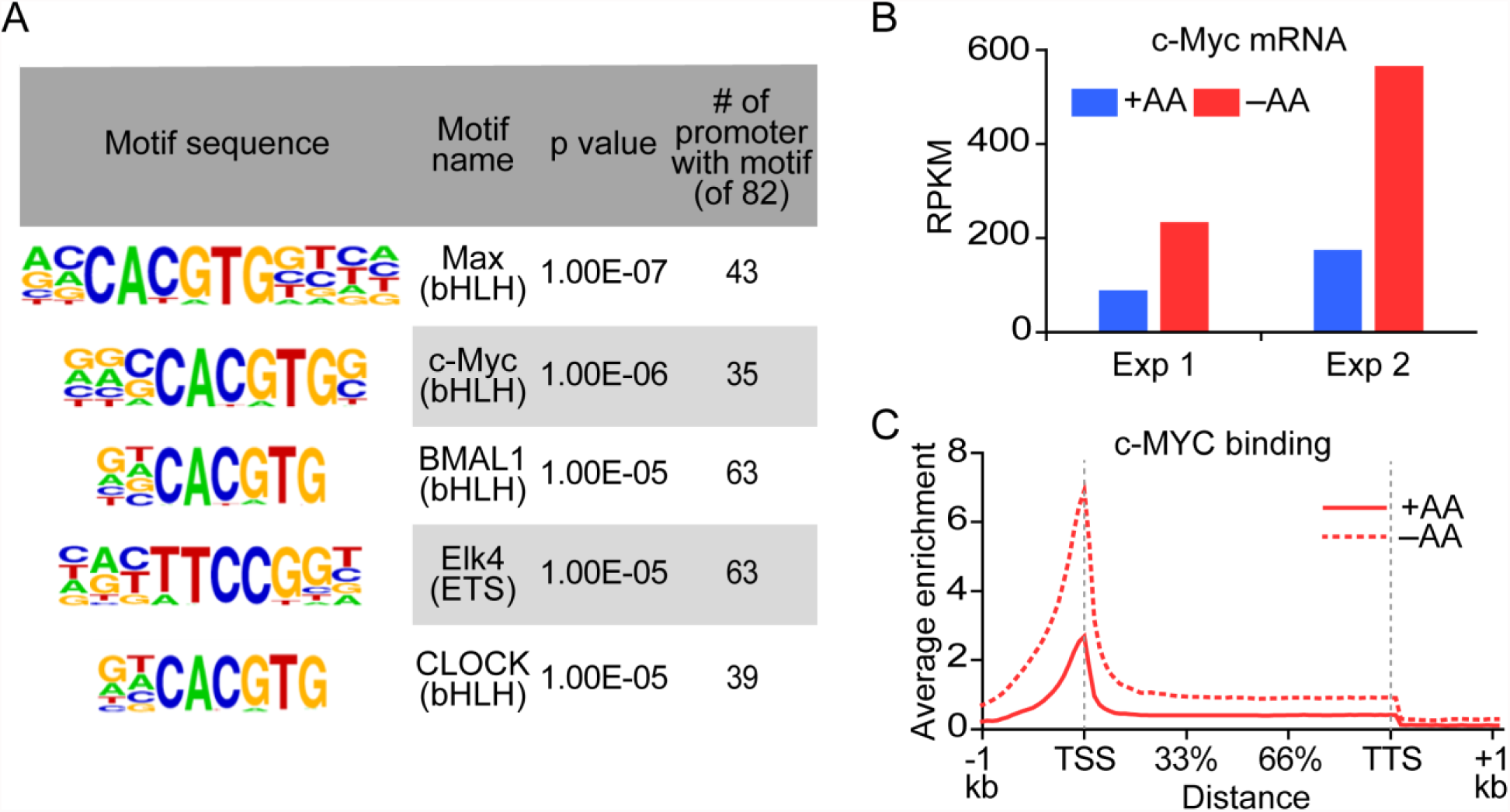
(A) Motif enrichment analysis of the promoters of upregulated genes involved in translational initiation and rRNA processing as shown in figure 2D. Regions from -500 to +750 bp of each promoter sequence were used for analysis. (B) Bar graphs showing upregulation of c-Myc mRNA in −AA compared to +AA from two independent RNA-seq experiments. (C) Metagene plot showing significant peaks of c-MYC binding in the same group of genes shown in figure 3A.

**Figure S4.**
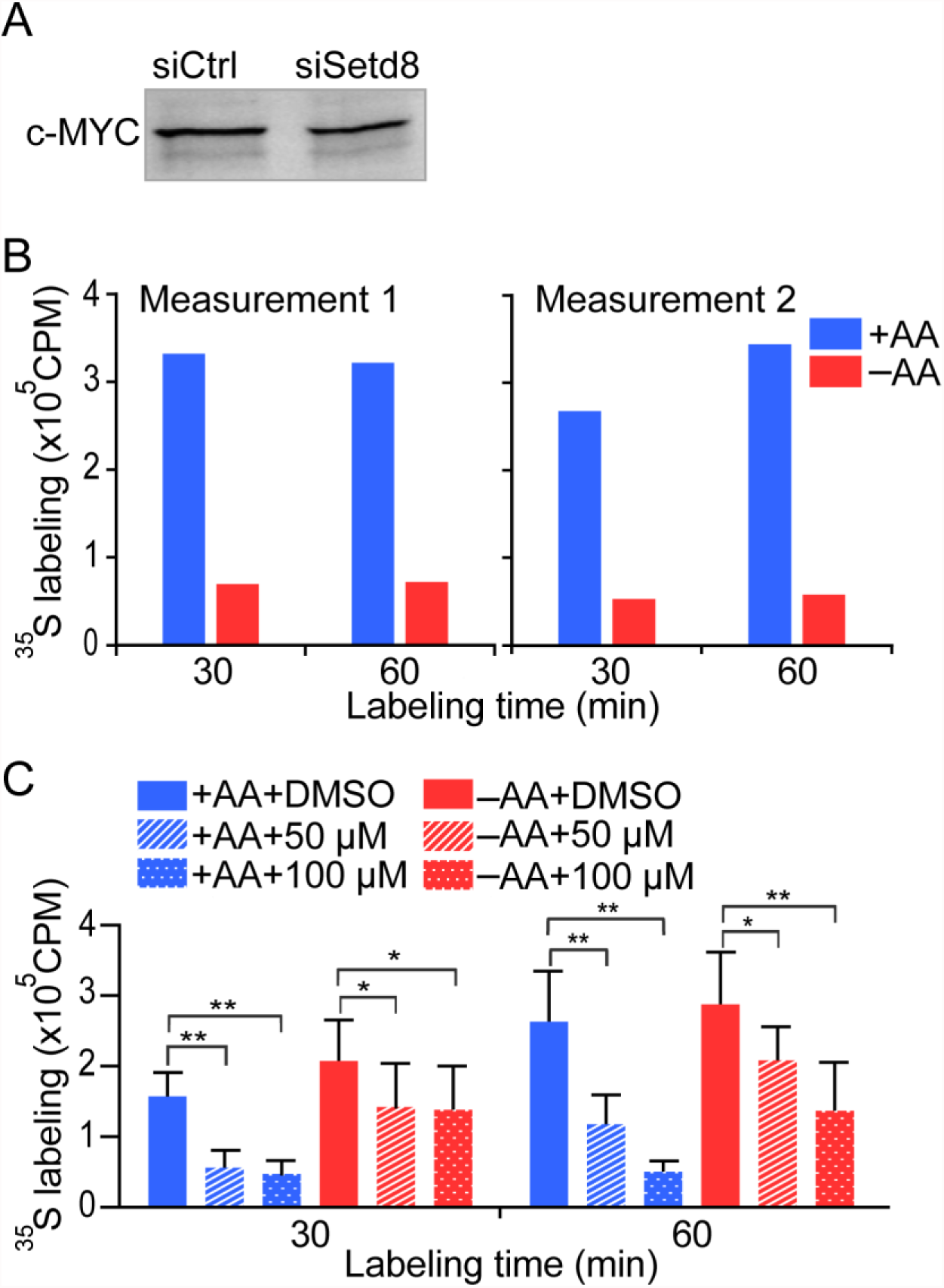
(A) WB of c-MYC in control and *SETD8* knockdown. (B) Levels of [^35^S]-methionine/cystiene incorporation by cells cultured in +AA or −AA for 16 hours prior to pulse labeling. In this experiment, —AA labeling medium contained only [^35^S]-methionine/cystiene with no additional amino acids to force the cells to rely on the internal pool of AAs for protein synthesis. (C) Primary data for figure 4E. Levels of [^35^S]-methionine/cystiene incorporation by cells treated with DMSO or indicated concentrations of c-MYC inhibitor 10058-F4 prior to pulse labeling. [^35^S]-methionine/cystiene labeling was done with the full complement of amino acids. *P<0.01, **P<0.0001.

